# Lipid–MOF Colloidosomes for Multimodal Encapsulation and Environmental Remediation

**DOI:** 10.64898/2026.03.24.714078

**Authors:** Jeshua Podliska, Rahul Dev Jana, Raheleh Ravanfar

**Author notes:** **Corresponding Author:** Raheleh Ravanfar, Assistant Professor, Department of Chemistry and Biochemistry, Texas Tech University, Lubbock, Texas 79409, United States.

## Abstract

The scalable fabrication of stable colloidosomes with controlled permeability and defined multicompartmental architecture remains a critical challenge, limiting their broader use in molecular delivery and environmental remediation. Here, we develop a hybrid lipid-metal-organic framework (lipid-MOF) colloidosome assembled through an interfacial emulsification strategy that integrates the structural rigidity of ZIF-8 particles with lipid-mediated membrane stabilization. During assembly, ZIF-8 particles accumulate at the oil-water interface to form a shell, producing hollow micron-sized spherical colloidosomes. The resulting colloidosomes exhibit excellent colloidal stability in aqueous media for over 30 days with a zeta potential of approximately −50 mV. Nitrogen adsorption measurements reveal a surface area of 45 m^2^g^-1^ and an average pore width of 4 nm. Fluorescence imaging shows that hydrophobic Nile red preferentially partitions into the colloidosomal membrane, whereas hydrophilic fluorescein isothiocyanate (FITC) localize predominantly within the aqueous interior, enabling simultaneous encapsulation of molecules with contrasting polarity with loading efficiencies approaching 90%. Furthermore, the colloidosomes demonstrate rapid removal of model pollutants from water, achieving >90% removal of methylene blue and metal ions without stirring. Together, these results introduce lipid–MOF colloidosomes as a new class of hybrid platforms that unify structural stability, multicompartmental encapsulation, and efficient adsorption behavior, opening pathways toward sustainable platforms for drug delivery and environmental bioremediation.

## INTRODUCTION

Colloidosomes, hollow microcapsules assembled from particles at emulsion interfaces, represent a versatile platform with tunable permeability, mechanical integrity, and chemical functionality.^1–6^ Their hollow architecture enables the confinement of functional species within a protected interior while the surrounding shell regulates transport of molecules across the membrane. As a result, colloidosomes have been explored applications ranging from controlled delivery and catalysis to sensing and environmental remediation.^2, 7–11^ Despite these promising opportunities, achieving colloidosomes that simultaneously exhibit structural robustness, controlled permeability, and multifunctional encapsulation remains a persistent challenge. In many particle-based colloidosomes, the shell is formed through the jamming of nanoparticles at fluid interfaces, often producing interparticle defects or voids that can compromise mechanical stability and limit precise control over molecular transport.^12^ MOFs have attracted significant attention as building blocks for advanced colloidal assemblies because of their exceptional high surface areas, tunable pore structures, and chemical versatility.^13, 14^ Zeolitic imidazolate framework-8 (ZIF-8) has been widely studied owing to its chemical stability, accessible microporosity, and compatibility with aqueous room-temperature synthesis conditions.^10, 15–21^ Recent studies have demonstrated that MOF can assemble at liquid interfaces to generate MOF-based colloidosomes and hollow capsules with potential applications in catalysis, pollutant capture, and self-propelled micromotors.^22–27^ While these approaches highlight the promise of MOF-derived colloidal shells, the resulting membranes typically consist of loosely packed nanocrystals, where interparticle gaps and structural heterogeneity may limit scalability, mechanical stability and reduce control over molecular partitioning within the capsule.^2, 28–33^ In biological systems, lipid membranes provide interfacial stability and polarity-dependent organization of molecules, while embedded proteins or inorganic cofactors introduce additional catalytic or transport functions. Inspired by these natural architectures, hybrid assemblies that combine lipid phases with inorganic materials can open new routes toward multifunctional compartmentalized systems.

Here, we engineered a cooperative lipid-MOF interfacial assembly strategy that enables the formation of lipid-MOF colloidosomes while preserving porous transport pathways across the shell. In this approach, ZIF-8 particles accumulate at the oil-water interface during emulsification to form a particulate shell, while a thermally stable lipid phase co-assembles at the interface. Rather than forming a continuous impermeable coating, the solidifying wax domains bridge adjacent MOF particles, sealing structural defects and reinforcing shell cohesion while maintaining accessible porous transport throughout the colloidosomal mesoscopic voids. This cooperative organization produces a mechanically stabilized yet permeable hybrid interface. Using high-shear homogenization, this wax-assisted interfacial assembly process provides a scalable route to monodisperse lipid-MOF colloidosomes with sustained colloidal stability in aqueous media. The resulting hierarchical architecture enables polarity-dependent spatial organization of guest molecules, where hydrophilic species localize preferentially within the aqueous interior and hydrophobic species partition into the lipid-integrated shell domain. Beyond molecular encapsulation, the aqueous interior functions as an accessible sequestration chamber for small organic molecules and metal ions. Small molecule pollutants diffuse through the porous composite shell and accumulate within the internal cavity, where the confined geometry and large internal surface area enhance uptake efficiency. This core-mediated capture mechanism distinguishes our system from motion-driven or purely surface-adsorption MOF assemblies. Together, these results establish lipid–MOF colloidosomes as a new class of hybrid colloidal materials that integrate structural reinforcement, multicompartmental encapsulation, and aqueous stability within a scalable platform for molecular delivery and environmental remediation.

## RESULTS AND DISCUSSION

While MOFs have been extensively employed as stabilizers in Pickering emulsions,^34, 35^ their cooperative assembly with a solid lipid phase to form structurally reinforced colloidosomes remains unexplored. We hypothesized that combining pre-formed MOF particles with a lipid matrix under double-emulsion conditions would enable the formation of hybrid colloidosomes, in which MOF particles assemble at the lipid–water interface and become embedded beneath a solidified lipid shell (Fig. 1a). This architecture could bridge interfacial defects and enhance the structural stability. To evaluate this concept, we apply a double-emulsion strategy in which an aqueous dispersion of pre-formed ZIF-8 was first emulsified within a molten lipid phase to generate a primary water-in-oil (W/O) emulsion, followed by secondary emulsification into an external aqueous stabilizer solution to produce water-in-oil-in-water (W/O/W) emulsions. Upon cooling, the lipid phase solidifies, yielding colloidosomal particles with a continuous outer lipid shell and MOF nanoparticles positioned at or near the internal interface. Moreover, the stabilizers used in the secondary phase can play an important role in stabilizing these lipid-MOF colloidosomes. Because interfacial and structural stability of the colloidosomes is affected by both lipid composition and surfactant type, we screened various lipid–surfactant combinations to identify formulations capable of maintaining long-term colloidal stability (Table S1).

**Figure 1.**
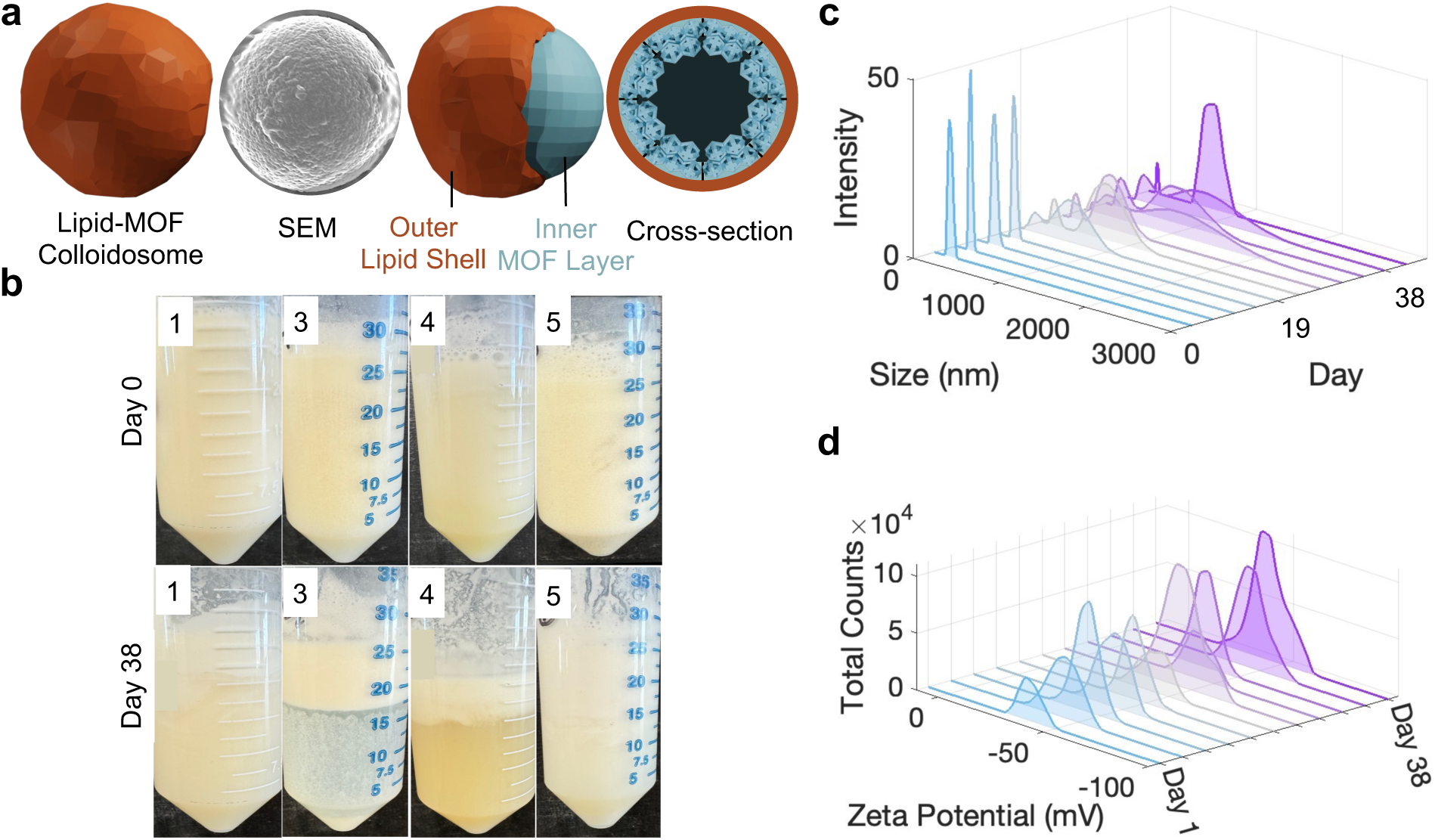
a) The schematic illustration of the assembly of lipid-MOF colloidosomes, demonstrating an external solid lipid shell with MOF particles decorated at the emulsion interface beneath the lipid layer. b) Representative images of different formulations, #1, #3, #4 and #5, showing their colloidal stability over 38 days (Table S1). Formulations #5, including Pluronic F-127 showed stability for 38 days. c) The hydrodynamic size for the formulation #5, including Pluronic F127, over 38 days. d) Zeta Potential for the formulation #5, including Pluronic F-127 over 38 days.

Formulations prepared with olive oil and stabilized using CTAB (formulations # 10-12) or guar gum (formulation # 3) as stabilizers in the external aqueous phase exhibited visible phase separation within 1-2 days, indicating instability of the colloidosomal structures and insufficient interfacial stabilization (Table 1). Given the practical advantages of natural waxes, including renewable origin, biocompatibility with food and biomedical contexts, and thermal robustness during processing,^36–42^ we next focused on wax-based lipid matrices. Coconut wax provided improved structural integrity during emulsification and resisted coalescence during storage (Fig. 1b, S1, and S2). When combined with Triton X-100 (formulation #1) or Pluronic F127 (formulation #5) as the external stabilizer, the formulation yielded homogeneous dispersions with polydispersity index (PDI) of 0.4 and 0.2, respectively, that remained colloidally stable for at least 38 days with no observable macroscopic phase separation (Figure 1b, S1, and S2). In this formulation, ZIF-8 MOFs were first synthesized in aqueous solution by mixing 2-methylimidazole with zinc nitrate under stirring at room temperature, yielding a milky dispersion. This ZIF-8 aqueous dispersion was added dropwise to a molten lipid phase composed of coconut wax, soy lecithin, and glycerol monooleate maintained at 70 °C under stirring, forming a primary water-in-oil (W/O) emulsion. This primary emulsion was subsequently introduced dropwise into an external aqueous solution of stabilizer containing Triton X-100, Pluronic F-127, or a combination of both under high-shear homogenization to generate a water-in-oil-in-water (W/O/W) double emulsion. Upon cooling to room temperature, the lipid phase solidified, producing colloidosomal particles in which ZIF-8 nanoparticles are organized at the interface beneath the solid lipid shell (Fig. 1a).

The colloidal stability was evaluated by monitoring the hydrodynamic diameter and zeta potential over a 38-day storage period (Figs. S1-S2). Dynamic light scattering (DLS) analysis show that the choice of stabilizer in the external aqueous phase significantly affects both the hydrodynamic diameter and polydispersity of the lipid-MOF colloidosomes (Figs. 1c-d and S1-S2). The coconut wax/Triton X-100 formulations yielded particle sizes in the 2–4 μm range (Fig. S1). Replacing Triton X-100 with Pluronic F-127 reduced the size of the lipid-MOF colloidosomes to 0.8 μm (Figs. 1c and S1-S2), which confirms the exceptional role of Pluronic F-127 in providing the hydrophilicity and steric stabilization compared to conventional nonionic surfactants.^43, 44^ Our results show that Pluronic F127 increases zeta potential of the lipid-MOF colloidosomes to −50 mV (Fig. 1d), providing high colloidal stability due to its triblock structure containing poly ethylene glycol block poly propylene glycol block poly ethylene glycol (PEG-PPG-PEG) with two hydrophilic polyethylene glycol chains. Our lipid-MOF colloidosomes demonstrated substantially enhanced stability compared to other reported ZIF-8 colloidosomes, which typically exhibit aggregation within 7-14 days.^45^ Formulation #5 was selected as the most stable system and was further improved with increasing the Pluronic F127 to 1% (formulation #6). This formulation was then used for all subsequent experiments described in this study.

Fourier-transform infrared (FTIR) spectra and elemental analysis confirmed the successful integration of ZIF-8 in the lipid-MOF colloidosomes (Figs. 2a, S3 and S4). The spectrum of lipid-MOF colloidosomes show the characteristic vibrational bands of ZIF-8, including the C=N stretching vibration of the imidazole ring at 1584 cm^-1^ and Zn–N coordination vibration at 421 cm^-1,33^ indicating that the MOF remains structurally intact during the lipid-MOF colloidosomal assembly (Fig. 2a). In addition, the lipid-MOF spectrum illustrates bands at 2920 cm^-1^ and 2850 cm^-1^ corresponding to the asymmetric and symmetric stretching modes of aliphatic hydrocarbon chains from the wax, confirming the presence of the lipid phase within the hybrid structure (Fig. 2a). Thermogravimetric analysis (TGA) was performed to evaluate the thermal profile of the lipid-MOF colloidosomes (Fig. 2b and S5). ZIF-8 shows a small initial mass loss below 200°C, followed by gradual decomposition of the imidazolate linkers at higher temperatures while retaining a significant inorganic residue at 800°C associated with zinc. In contrast, the lipid-MOF colloidosomes exhibit a substantial mass loss beginning at 250°C and progressing rapidly between 300°C and 420°C, consistent with thermal degradation of the lipid component (Fig. 2b and S5). The significantly larger mass loss observed for lipid-MOF colloidosomes relative to ZIF-8 alone reflects both the dilution of the inorganic framework by the lipid phase and the modified decomposition pathway of lipid-MOF colloidosomes during thermal treatment. Scanning electron microscopy (SEM) images revealed the formation of spherical lipid-MOF colloidosomes with diameters in the 1-2 μm range (Fig. 2c-f). The smooth and continuous outer surface of the lipid-MOF colloidosomes indicates successful encapsulation of the MOF phase beneath the lipid shell (Fig. 2c-f). Higher magnification SEM images show partially fractured colloidosomes, which provide visual evidence of a hollow interior architecture formed through emulsion templating (Fig. 2e-f). In figure 2f, contrast variations within the particle shell suggest the presence of a denser MOF-rich region beneath the outer lipid layer, while the darker central region is consistent with an empty interior volume after solvent removal during sample preparation. This morphology differs significantly from conventional MOF colloidosomes assembled solely from aggregated crystalline MOFs, which typically exhibit rough, granular shells with visible interparticle voids.^19,46^ Instead, the lipid-MOF colloidosomes display smoother and more cohesive shell structure (Fig. 2c-f). This behavior is attributed to lipid matrix which fills interstitial spaces between ZIF-8 particles and produces a more continuous composite shell.

**Figure 2.**
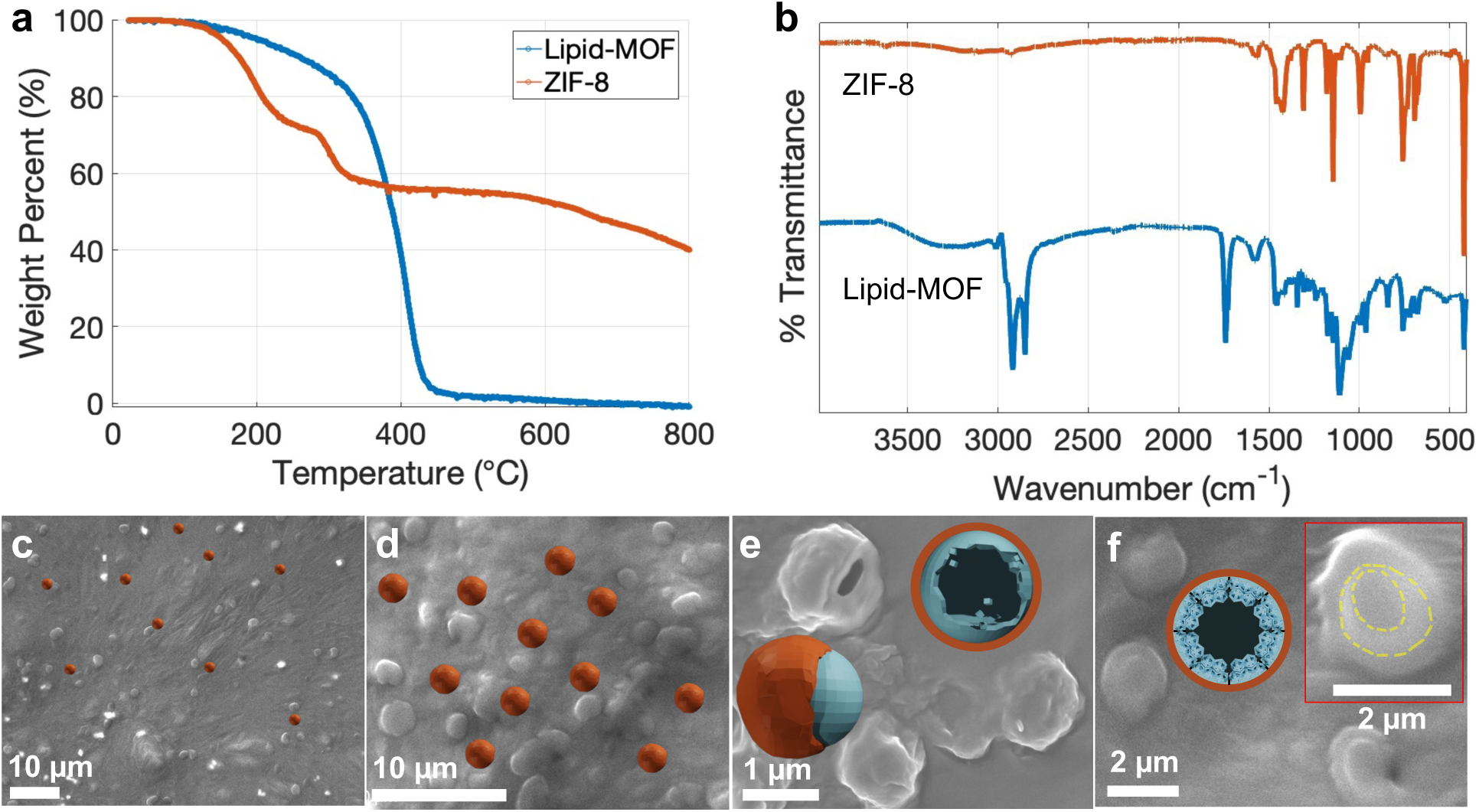
a) FTIR spectra for lipid-MOF colloidosomes and ZIF-8. b) TGA for the lipid-MOF colloidosomes and ZIF-8. c) SEM image of the lipid-MOF colloidosomes appeared well-dispersed. d) Magnification SEM images of the lipid-MOF particles. e) The fractured lipid-MOF colloidosome with a hollow interior domain. f) Contrast variations within the lipid-MOF colloidosomes, illustrating the surface lipid shell, MOF layer, and hollow inner domain.

Previous studies on encapsulation have largely focused on either hydrophilic or hydrophobic cargo independently, which was limited to single-polarity systems.^47–57^ Combining both types of molecules within a single carrier, however, remains a significant challenge due to their fundamentally different physicochemical properties. An important feature of the lipid–MOF colloidosomes is their ability to spatially compartmentalize molecules according to their polarity, enabling the coexistence of hydrophobic and hydrophilic species within distinct domains of the same carrier. To evaluate this compartmentalization behavior, we investigated the encapsulation of fluorescent probes with contrasting polarity using fluorescein isothiocyanate (FITC) as a hydrophilic model compound and Nile Red (NR) as a hydrophobic probe. Confocal laser scanning microscopy (CLSM) revealed clearly differentiated localization patterns for the two dyes (Fig. 3a). Nile Red exhibited strong red fluorescence predominantly at the lipid–MOF colloidosomal shell due to its lipophilic nature (Fig. 3a). In contrast, FITC displayed yellow fluorescence primarily within the interior of the particles, indicating accumulation in the aqueous cavity of the colloidosomes (Fig. 3a). Overlay images further confirm this spatial separation, showing a bright ring red emission surrounding the yellow interior. These observations demonstrate that the hybrid architecture establishes chemically distinct domains, a hydrophobic lipid-MOF interfacial shell and a hydrophilic internal cavity, allowing selective localization of molecules according to their polarity (Fig. 3a). Such spatial segregation within a single microcarrier highlights the potential of lipid-MOF colloidosomes as versatile platforms for co-encapsulation, controlled release, and multifunctional systems.

**Figure 3.**
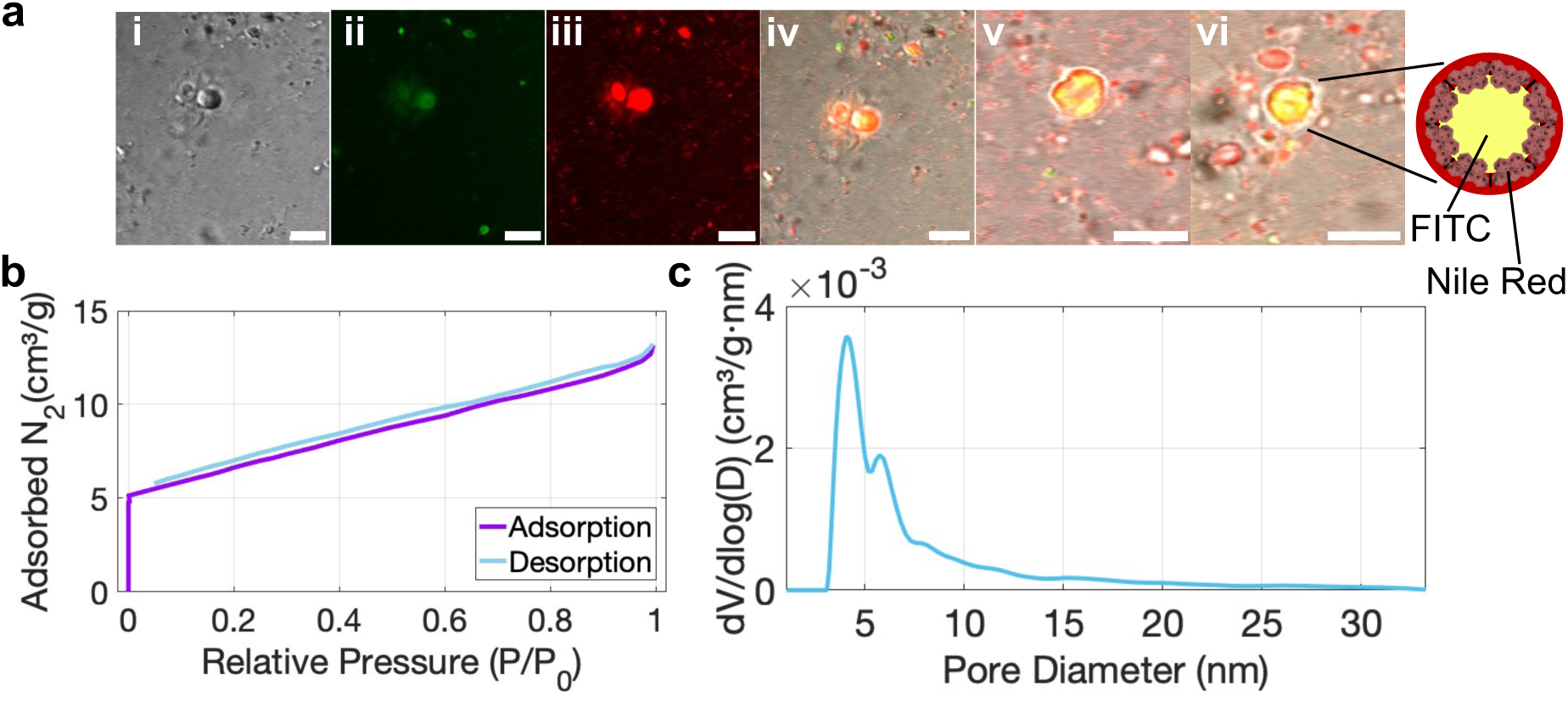
a) Confocal laser scanning microscopy images of lipid-MOF colloidosomes, i: bright field, ii: excited at 488 nm, iii: excited at 535 nm, iv, v & vi: overlay images show the lipid-MOF colloidosomes with red shell and yellow core due to the accumulation of Nile red and FITC, respectively; Scale bars: 10 μm. b) Nitrogen adsorption and desorption curves. c) Density functional theory (DFT) pore size distribution detected with N_2_ adsorption and desorption at 77 K for lipid-MOF colloidosomes.

Nitrogen adsorption measurements revealed a Brunauer–Emmett–Teller (BET) surface area of 46 mg^-1^ and a pore volume of 0.02 cc g^-1^ for the lipid-MOF colloidosomes (Figs. 3b, S6, and Table S2), substantially lower than values typically reported for pristine ZIF-8. This reduction reflects the presence of lipid domains that partially cover the external surfaces of the MOF nanocrystals and fill interparticle voids within the colloidosome shell, thereby limiting nitrogen accessibility under cryogenic adsorption conditions. The average pore diameter obtained from DFT analysis (∼4 nm) suggests that the accessible porosity is dominated by mesoporous domains arising from interparticle spaces rather than the intrinsic microporosity of ZIF-8 (Fig. 3c and Table S2).

Wastewater generated from industrial processes rarely contains a single type of contaminant; instead, it typically consists of complex mixtures of organic dyes, dissolved metal ions, and other chemical species. Despite this reality, most remediation studies still rely on simplified model systems that examine the removal of only one pollutant class at a time. The release of synthetic dyes into natural water systems is a particularly pressing environmental concern because these compounds are often chemically stable, highly visible even at low concentrations, and potentially harmful to both ecosystems and human health.^58, 59^ Methylene blue (MB), for example, is widely used in industrial dyeing and biological staining processes, yet its accumulation in water sources has been associated with adverse health effects including respiratory irritation, gastrointestinal distress, and eye irritation.^60–62^ Materials capable of addressing multiple contaminant types simultaneously would therefore represent a meaningful step toward more practical wastewater treatment technologies. In the present work, we explored whether lipid–MOF colloidosomes could serve as versatile adsorption platforms capable of removing both organic dyes and metal ions from aqueous environments. Given the hierarchical structure and amphiphilic interfaces present in our lipid–MOF colloidosomes, we reasoned that these assemblies might provide an efficient platform for dye capture. Indeed, when exposed to aqueous MB solutions, the colloidosomes demonstrated near-complete removal of the dye without stirring (Fig. 4a). Dynamic light scattering measurements revealed a clear increase in particle size after dye exposure, reaching approximately 4 μm, which suggests that MB molecules are not simply adhering to the outer surface but are being taken up into the hollow interior of the assemblies (Fig. S7). Moreover, the adsorption without stirring was more efficient than with stirring (Fig. S8). This observation is consistent with a mechanism in which the porous MOF framework and lipid interface work together to concentrate hydrophilic dye molecules within the internal cavity of the colloidosomes. Moreover, the encapsulation efficiency of MB in the lipid-MOF colloidosomes was over 90%, which shows the intriguing capability of these colloidosomes for entrapping the hydrophilic molecules (Fig. 4a).

**Figure 4.**
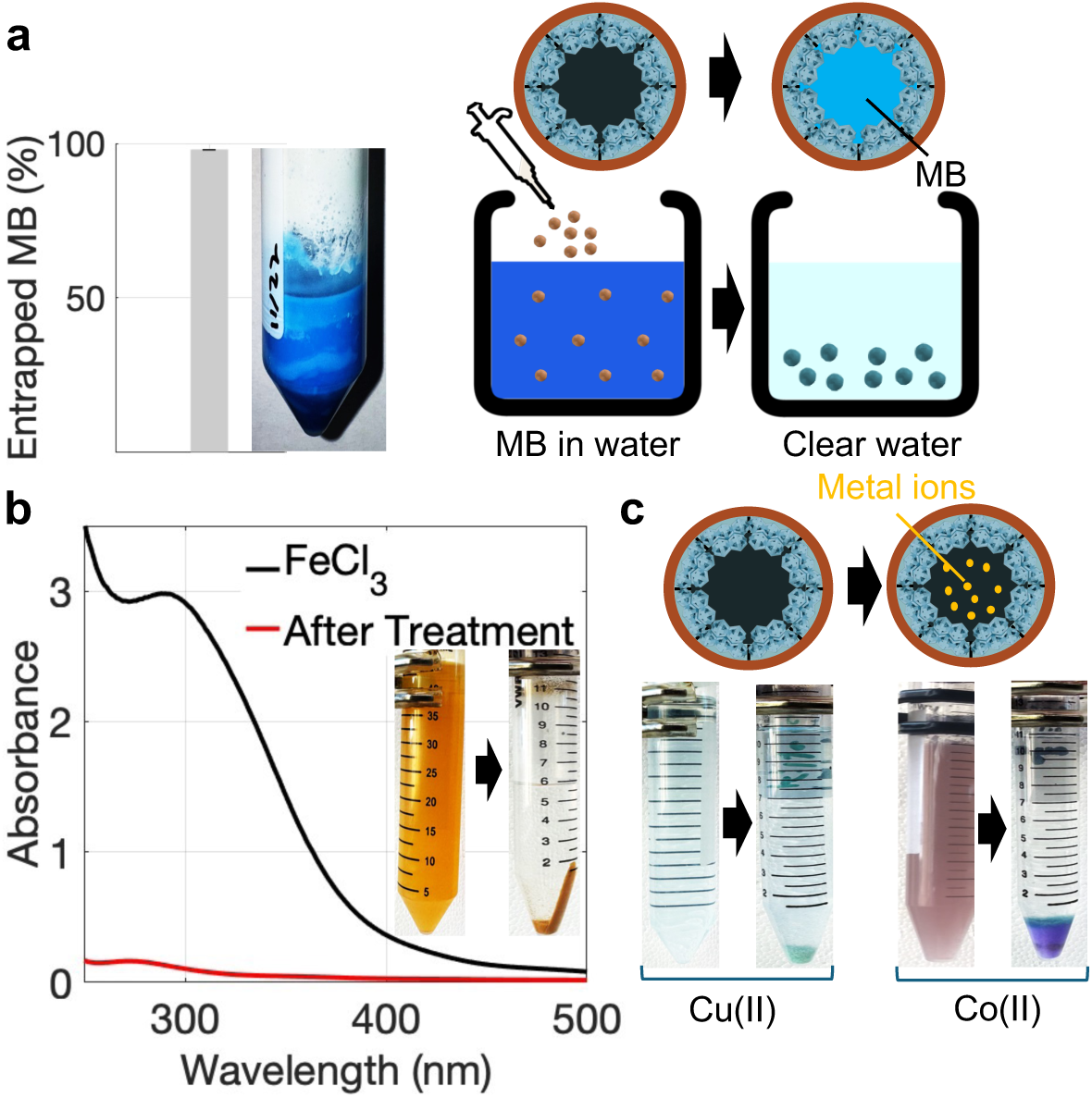
a) Encapsulation efficiency of MB and the removal performance of the lipid-MOF colloidosomes. b) UV-vis spectra and optical images illustrating the Fe(III) adsorption capacity of lipid-MOF colloidosomes without stirring. c) Optical images of Cu(II) and Co(II) adsorption capacity of lipid-MOF colloidosomes.

Heavy metal contamination represents another major environmental challenge associated with industrial activity. Metals such as mercury, cadmium, lead, chromium, and arsenic can persist in aquatic systems and pose serious risks due to their toxicity and ability to bioaccumulate.^63^ To explore whether the lipid-MOF colloidosomes could also address this class of pollutants, we investigated their ability to capture dissolved metal ions using Fe(III), Cu(II), and Co(II) as representative examples. Upon introduction of the colloidosomes into a solution containing FeCl_3_ (1000 ppm), or CoSO_4_ (100 ppm) or CuCl_2_ (100 ppm), the lipid-MOF colloidosomes rapidly interacted with the metal-containing environment without stirring, accompanied by a visible color change that indicated metal uptake (Fig. 4b-c, Supplementary videos 1-3). The lipid-MOF colloidosomes gradually aggregated and settled to the bottom of the solution after the adsorption process was complete (Fig. 4b-c, Supplementary videos 1-3). Quantitative UV–vis analysis confirmed that more than 90% of the iron, cobalt, and copper were removed from solution (Figs. 4b-c, and S9-S10), suggesting that the reduced BET surface area does not preclude efficient adsorption in aqueous environments. Instead, the hybrid architecture of lipid-MOF colloidosomes introduces interfacial adsorption sites and mesostructured pathways that facilitate the capture of small organic molecules and metal ions. This platform may provide a promising starting point for the development of colloidal assemblies capable of capturing a broader range of heavy metal contaminants and organic dyes from wastewater.

## CONCLUSION

In this work, we developed hybrid lipid-MOF colloidosomes through an interfacial emulsification strategy that brings together the structural robustness of ZIF-8 with the stabilizing properties of a solid lipid phase. This cooperative assembly produces hollow colloidosomes with diameters of approximately 1–2 μm that remain colloidally stable in aqueous environments for extended periods. The architecture of these colloidosomes enables spatial organization of molecules according to their polarity. Confocal fluorescence imaging shows that hydrophobic Nile Red preferentially associates with the lipid-rich shell, while hydrophilic FITC accumulates within the aqueous interior. This behavior highlights the ability of the hybrid structure to accommodate molecules with different chemical characteristics within separate domains of the same particle. Nitrogen adsorption measurements further suggest that the accessible porosity of the system is largely associated with mesoscopic interparticle spaces within the colloidosomal shell. Beyond encapsulation, the lipid-MOF colloidosomes also exhibit promising adsorption behavior toward aqueous contaminants. The particles efficiently removed more than 90% of methylene blue dye and representative metal ions such as Fe(III), Cu(II), and Co(II) from solution without the need for stirring. These observations indicate that the hollow interior and porous shell together provide an effective environment for capturing dissolved species from water. Overall, these results demonstrate that combining lipid phases with MOF particles at emulsion interfaces offers a simple and scalable route to multifunctional colloidal assemblies. The resulting lipid-MOF colloidosomes integrate structural stability, compartmentalized encapsulation, and pollutant capture within a single platform. Such hybrid architectures may provide useful starting points for future developments in areas ranging from controlled delivery systems to environmental remediation technologies.

## MATERIALS AND METHODS

### Materials

Pluronic F-127, 2-methylimidazole, Span® 80, hexadecyltrimethylammonium bromide (CTAB), copper (II) chloride, iron (III) chloride, and cobalt (II) sulfate heptahydrate were obtained from Sigma-Aldrich, MO, USA. Triton X-100, fluorescein isothiocyanate, Nile red, methylene Bue, and palmitic acid were obtained from Thermo Fisher Scientific, MA, USA. Zinc nitrate hexahydrate and nickel sulfate were obtained from Ward’s Science, ON, CAN. Coconut wax was obtained from American Soy Organics IW, USA. Liquid soy lecithin was obtained from Velona, IL, USA. Glycerol monooleate was obtained from Spectrum, NJ, USA. Pure olive oil was obtained from Albertsons Companies, ID, USA.

### Preparation of Lipid-MOF Colloidosomes

Briefly, 1 mL of 0.1 M Zinc Nitrate was added to 1 mL of 2.5 M 2-methylimidazole while stirring at 1000 rpm for 20 minutes at room temperature to form a milky white dispersion of ZIF-8 MOF. The lipid phase, containing coconut Wax (1.2 g), soy lecithin (0.35 g) and glycerol monooleate (GMO) (0.35 g) was heated to 70°C and stirred at 1000 rpm for 10 minutes. The preformed MOF solution was then added dropwise to the lipid phase and left to stir for 10 minutes forming the W/O emulsion. The resulting emulsion was diluted to an aqueous phase consisting triton X-100 (1% w/w), or Pluronic F-127 (0.5% w/w) under high shear homogenization at 15000 rpm for 15 minutes to generate W/O/W double emulsion while the lipid is solidified at room temperature to form the lipid shell, where the ZIF-8 particles are oriented beneath the lipid shell at the emulsion interface.

### Confocal Laser Scanning Microscopy (CLSM)

To investigate polarity-dependent partitioning within the lipid–MOF colloidosomes, Nile Red and fluorescein isothiocyanate (FITC) were used as fluorescent probes for hydrophobic and hydrophilic domains, respectively. Nile Red preferentially stains the lipid-rich shell, whereas FITC localizes within the internal aqueous cavity, enabling visualization of the spatial organization of hydrophobic and hydrophilic regions within the colloidosomal architecture. Confocal imaging was performed using an Olympus FV3000 laser-scanning confocal microscope (Olympus Corp., Tokyo, Japan) equipped with 100× objective lenses. Fluorescence images were acquired using FITC and propidium iodide (PI) excitation and emission filter sets, along with differential interference contrast (DIC) imaging.

### Dynamic Light Scattering (DLS)

Particle size distribution of Lipid-MOF colloidosomes was monitored using Malvern Zetasizer Pro (Malvern Panalytical Ltd., UK) in different formulations. The colloidal stability of the particles was assessed through zeta potential measurements with an electroactive cell (DS1070). Samples were tested for 3 times to record average particle size and polydispersity index (PDI).

### Thermogravimetric Analysis (TGA)

Thermal stability of lipid-MOF colloidosomes was assessed using TGA (Shimadzu DTG-60H, Shimadzu Corp., Kyoto, Japan). Emulsions were centrifuged at 20,000 ×g for 30 minutes and the pellet was dried using SpeedVac (Thermo Fisher Savant Speed Vac Plus SC110A, Savant Instruments Inc, US). The dried samples were placed in alumina crucibles (AL6211 6 mm) and heated from room temperature to 800°C under a nitrogen atmosphere at a heating rate of 10 °C min^-1^.

### Fourier Transform Infrared Spectroscopy (FTIR)

Chemical Composition of emulsions were analyzed using ATR-FTIR (Nicolett FTIR, ThermoFisher Scientific, Thermo Electron Scientific Instruments LLC, USA).

### Ultraviolet–visible (UV-vis) Spectrophotometry

UV-vis spectrophotometry was conducted in the range of 190-900 nm, with 1.0 nm data interval, and 0.004 s averaging time on an Agilent Cary UV-Vis Compact Peltier from Agilent Technologies, Mulgrave, Australia.

### Scanning Electron Microscopy (SEM)

Surface morphology was investigated by scanning Electron Microscopy (SEM) using a Zeiss crossbeam 540 FIB-SEM equipped with Oxford energy-dispersive X-ray spectroscopy (EDS) system from ZEISS, 07745 Jena, Germany for elemental analysis. The SEM images were collected at 5 kV accelerating voltage and 5.9 mm working distance with side mount secondary electron detector. Samples were freeze dried using a Labconco Freezone (Labconco Corp., Kansas City, MO, USA) for 48 hours, placed on aluminum stubs covered with carbon adhesive tape, and sputter-coated with iridium using a Technics Hummer V Sputter Coater (Anatech USA, Union City, CA, USA).

### Brunauer–Emmett–Teller (BET) Analysis

Surface area analysis was performed using N_2_ adsorption-desorption isotherms at 77K with a Quantachrome® ASiQwin™ analyzer (Quantachrome Instruments, US). Prior to measurements, samples were degassed under vacuum at 100°C for 24h to eliminate adsorbed gases and moisture. The specific surface area of the samples was determined using the Brunauer–Emmett–Teller (BET) method. The BET equation was applied in the relative pressure range (P/P₀) of 0.05–0.50, where the isotherms showed linearity. The surface area was calculated based on the measured adsorption data. Pore size distribution and pore volume were further analyzed using Density Functional Theory (DFT) applied to the N_2_ adsorption-desorption isotherms. The analysis was carried out using N_2_ at 77 K on carbon (cylindrical pores, QSDFT adsorption branch). Model selection was based on the expected pore shape determined from SEM.

### Encapsulation Efficiency of Small Molecules in Lipid-MOF Colloidosomes

In order to investigate the capability of lipid-MOF colloidosomes for encapsulating the hydrophilic small molecules, we encapsulate methylene blue as a template. Briefly, 0.05 g MB was dissolved in the primary aqueous phase containing the precursors of ZIF-8 and applied in the formation of double emulsions as explained in the previous section. The lipid-MOF colloidosomes were then centrifuged and the MB was measured at 664 nm using M.B standard curve in the supernatant using Cary 60 UV-vis spectrophotometry (Agilent Technologies, USA). This amount was subtracted from the initial MB amount to calculate the encapsulation efficiency of MB in the lipid-MOF colloidosomes.

### Adsorption of Small Molecules using Lipid-MOF Colloidosomes

The best adsorption efficiency for methylene blue was associated to the sample that was synthesized using 10 mL 0.1 M Zinc nitrate, 10 mL of 2.5 M 2-methylimidazole, 3% w/w Pluronic F-127, 0.6 g coconut wax, and 0.15 g soy lecithin and GMO. Briefly, 1 mL sample was added to 2, 4, and 8 mL solution of 500 ppm methylene blue for 1 hour. The samples were then centrifuged at 20,000 ×g and the supernatant was analyzed by UV-vis spectrophotometry to monitor the residual of MB and used to calculate the adsorption capacity of lipid-MOF colloidosomes with and without stirring.

### Adsorption of Metal Ions using Lipid-MOF Colloidosomes

The best adsorption efficiency for metal ions was associated to the sample that was synthesized by preparing the lipid phase mixed with hexane and Span 80, and using 3 mL of 0.1 M zinc nitrate and 3 mL 2.4 M 2-methylimidazole. Briefly, 1 mL of this sample was added to 6 mL aqueous solution of iron(III) chloride (1000 ppm), or Copper(II) Chloride (100 ppm), or Cobalt(II) Sulfate (100 ppm) and incubated for 1 hour. Then, the samples were centrifuged and the supernatant was monitored for the trace of metal ions using UV-vis spectrophotometry.

## ASSOCIATED CONTENT

**Supporting Information**

The Supporting Information is available free of charge on the ACS Publications website.

Details on characterizations of lipid-MOF colloidosomes, data analysis using MATLAB, Figures S1–S10, Tables S1–S2 (PDF).

## AUTHOR INFORMATION

**Authors**

**Jeshua Podliska**, Department of Chemistry and Biochemistry, Texas Tech University, Lubbock, Texas 79409, United States

**Rahul Dev Jana**, Department of Chemistry and Biochemistry, Texas Tech University, Lubbock, Texas 79409, United States

## Author Contributions

The manuscript was written through contributions of all authors. All authors have given approval to the final version of the manuscript.

## Notes

The authors declare no competing financial interest.

## ACKNOWLEDGMENTS

The authors thank Bo Zhao, Anthony Cozzolino, and Juliusz Warzywoda for their assistance with data acquisition for SEM, FTIR, TGA, BET, and elemental analysis. We acknowledge support from the College of Arts & Sciences Microscopy. We are grateful for the funding support from the ACS Petroleum Research Fund Award 69034-DNI3.

## Data Availability Statement

The supplementary information, raw data and custom analysis scripts are available upon request from the corresponding author.

