## Supplementary material for "Lipid–MOF Colloidosomes for Multimodal Encapsulation and Environmental Remediation": SI

Table S1. Optimization of various formulations to obtain the most colloidal stability.

| **Formulas** | **Oil phase** | **Surfactant in Oil Phase** | **Surfactant in Aqueous Phase** |
|---|---|---|---|
| 1 | Coconut Wax | GMO + Soy Lecithin | Triton X-100 (0.5%) |
| 2 | Coconut Wax | GMO + Soy Lecithin | Triton X-100 (1%) |
| 3 | Coconut Wax | GMO + Soy Lecithin | Triton X-100 + Guar Gum |
| 4 | Bees Wax | GMO + Soy Lecithin | Triton X-100 |
| 5 | Coconut Wax | GMO + Soy Lecithin | Pluronic acid (0.5%) |
| 6 | Coconut Wax | GMO + Soy Lecithin | Pluronic acid (1%) |
| 7 | Palmitic Acid | GMO + Soy Lecithin | Triton X-100 |
| 8 | Coconut Wax | GMO + Soy Lecithin | Triton X-100 + Alginate |
| 9 | Palmitic Acid | GMO + Soy Lecithin | Triton X-100 + Alginate |
| 10 | Olive Oil | GMO | CTAB |
| 11 | Olive Oil | GMO + Soy Lecithin | CTAB |
| 12 | Olive Oil | Soy Lecithin | CTAB |
| 13 | Hexane | Span® 80 | Pluronic acid |

Table S2. Surface area, pore width, and pore volume of lipid-MOF colloidosomes derived from multipoint Brunauer–Emmett–Teller (BET) and Density Functional Theory (DFT) analysis.

| | |
|---|---|
| Pore Volume | 0.018 cc/g |
| Surface area | 45.9 $m^2/g$ |
| Lower confidence limit | 1.066 nm |
| Fitting Error | 1.503 % |
| Average Pore Width | 4.1 nm |

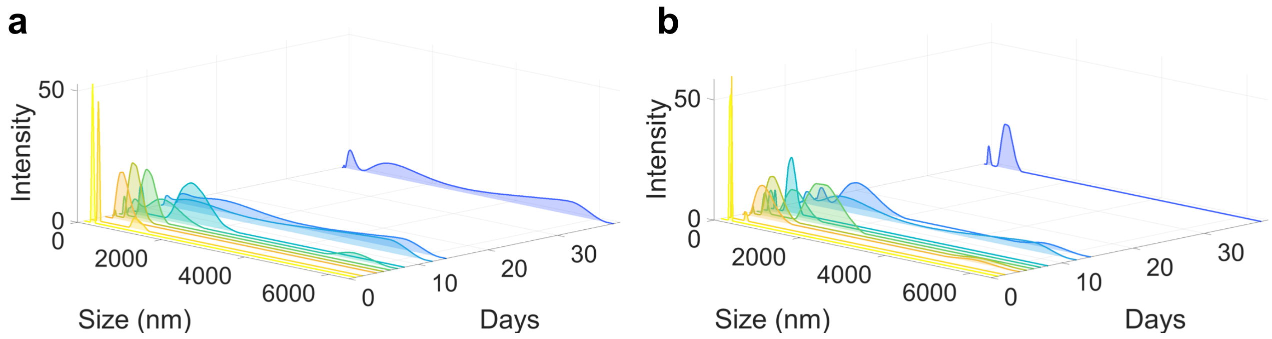


Figure S1. Particle size measurements over 38 days for formulations a) #5, and b) # 2.

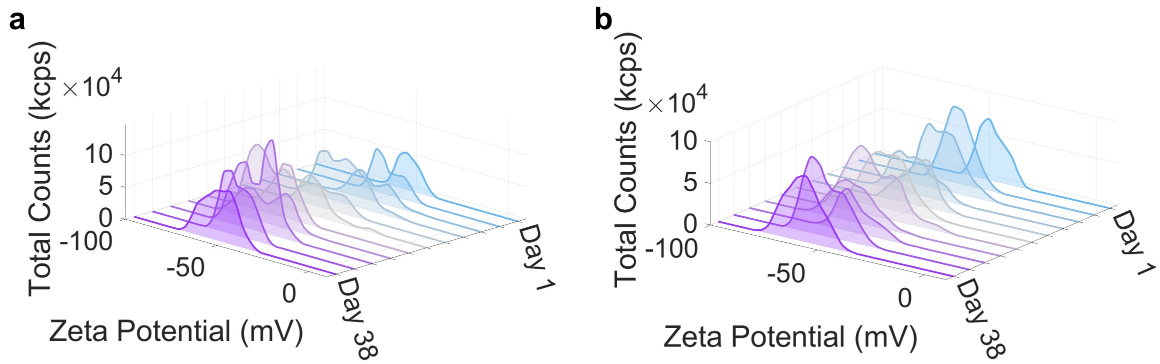


Figure S2. Zeta Potential measurements over 38 days for formulations a) # 1, b) # 5.

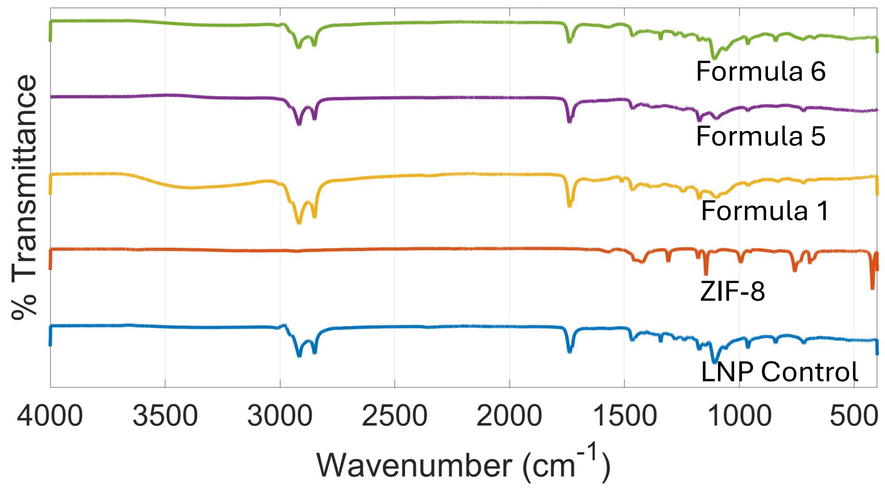


Figure S3. FTIR spectra for lipid-MOF colloidosomes synthesized through formulations #1, #5, and #6, ZIF-8, and lipid nanoparticles (LNP control).

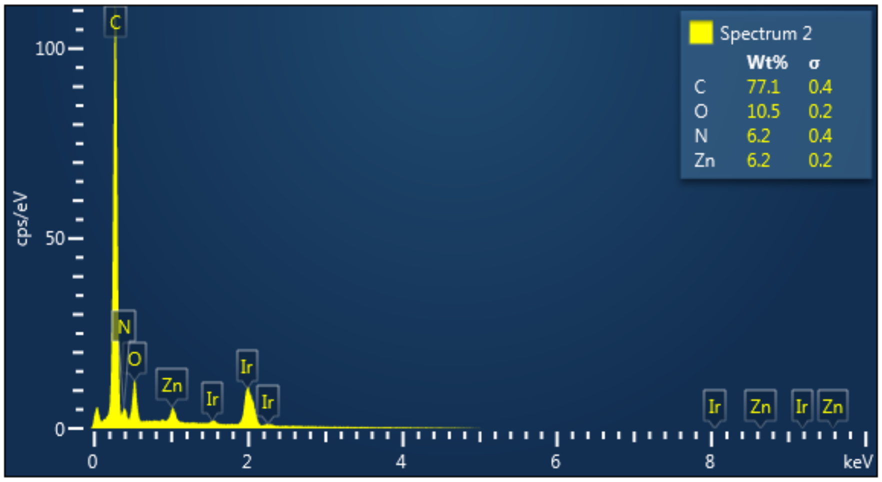


Figure S4. Elemental analysis confirms the presence of zinc in the lipid-MOF colloidosomes.

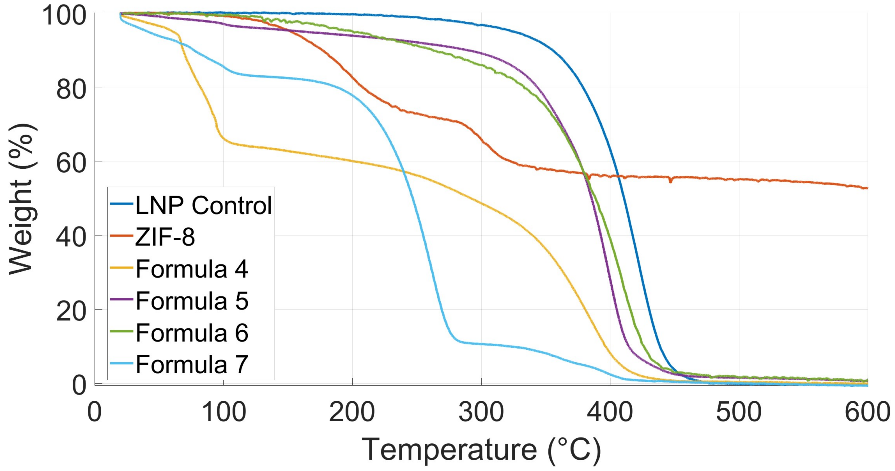


Figure S5. TGA for the lipid-MOF colloidosomes synthesized through formulations #4, #5, #6, #7, ZIF-8, and lipid nanoparticles (LNP control).

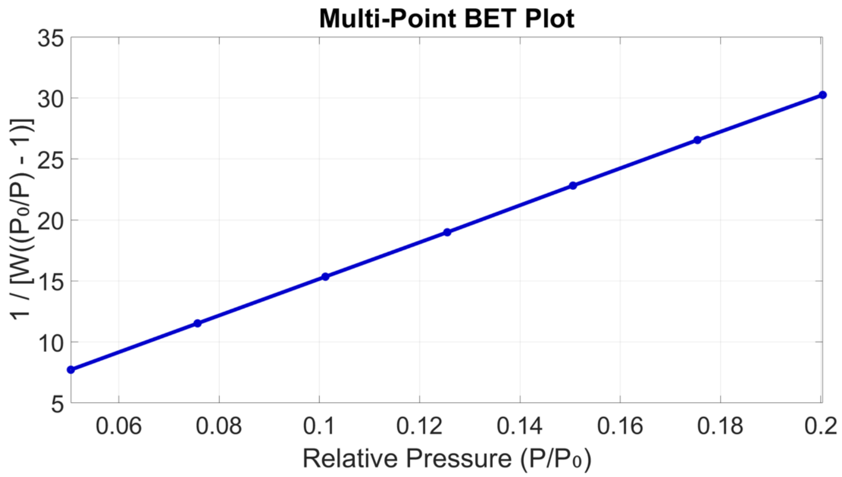


Figure S6. Multi-point BET plot of pore size distribution, showing that Pluronic F-127 carries a distinct mesoporous structure.

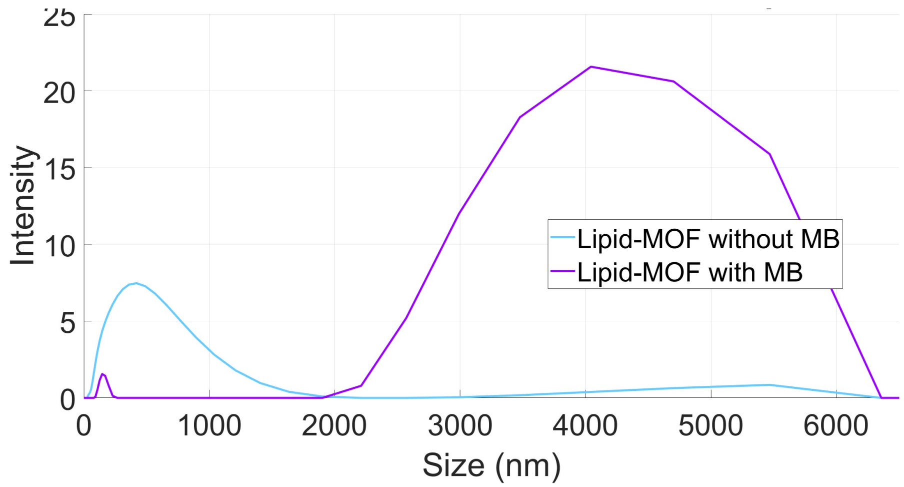


Figure S7. Increasing the size of lipid-MOF samples after adsorption of MB.

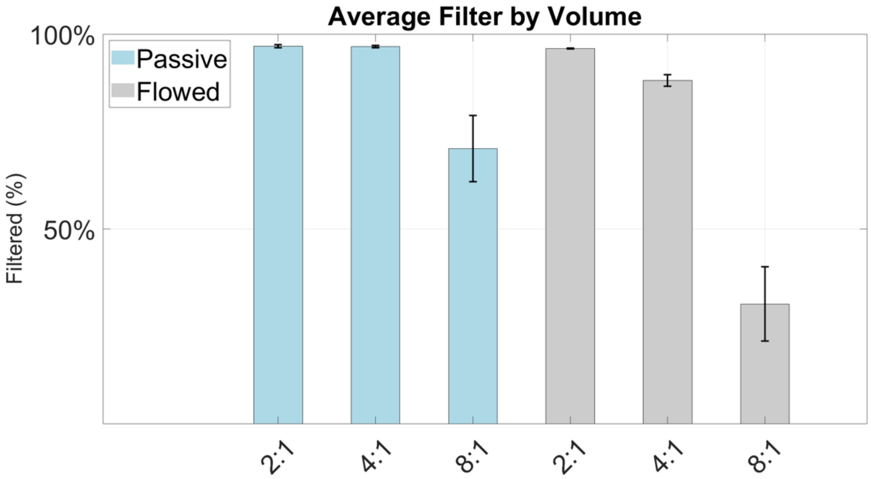


Figure S8. Removal of MB from 2, 4, and 8 mL solution of 500 ppm methylene blue using 1 mL of lipid-MOF colloidosomes after 1 hour with stirring (gray color, flowed) and without stirring (blue color, passive).

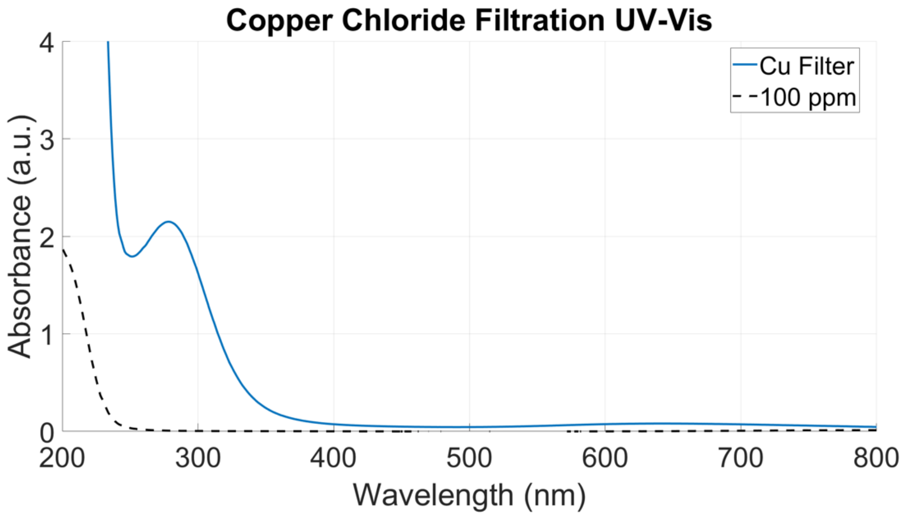


Figure S9. UV-vis spectra illustrating the Cu(II) adsorption capacity of lipid-MOF colloidosomes without stirring.

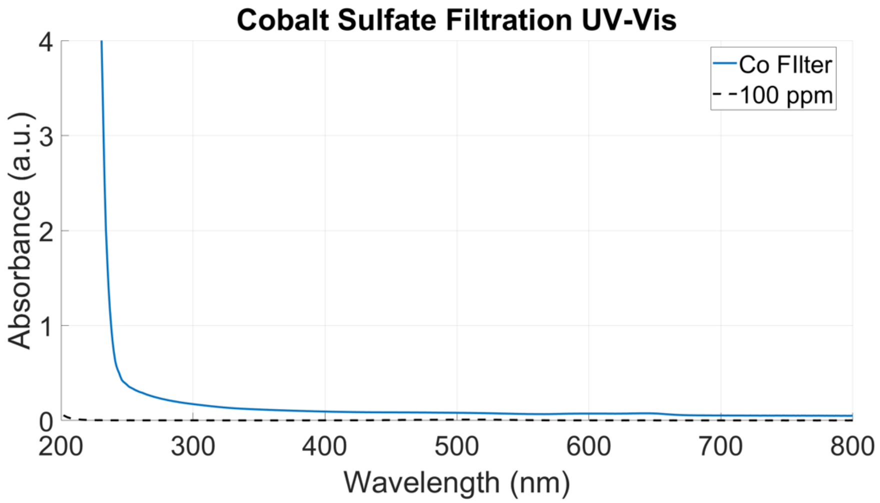


Figure S10. UV-vis spectra illustrating the Co(II) adsorption capacity of lipid-MOF colloidosomes without stirring.
